# Preference-Based Fine-Tuning of Genomic Sequence Models for Personal Expression Prediction with Data Augmentation

**DOI:** 10.1101/2025.11.09.687505

**Authors:** Moonwon Choi, Bokeum Cho, Seunggeun Lee

## Abstract

Despite substantial progress in genomic foundation models, accurately predicting inter-individual variation in gene expression from DNA sequence alone remains a major challenge. Current sequence-based models, such as Enformer and Borzoi, trained exclusively on the reference genome, cannot capture the effects of individual-specific regulatory variants. Moreover, the acquisition of paired whole-genome and transcriptome data required for personalized modeling is hindered by privacy and data-sharing constraints. To address this limitation, we integrate genomic data synthesis with established statistical frameworks. Our approach generates thousands of synthetic training samples by simulating genetic variation from the 1000 Genomes Project and assigning pseudo-expression labels using PrediXcan, a validated eQTL-based predictor. Because simulated and real expression values differ in scale and distribution, we introduce a preference-based objective that models relative rather than absolute expression patterns. Fine-tuning Enformer through alternating cycles of real-data regression and synthetic-data preference optimization enables efficient learning from both real and synthesized data. Using the GEUVADIS dataset, our framework outperforms AlphaGenome, PrediXcan, and Enformer fine-tuned without synthesized data, demonstrating that simulation-based integration of population-level regulatory knowledge can effectively mitigate data scarcity and improve cross-individual generalization in sequence-based gene expression prediction.

**Availability and implementation:** Code and data are available at https://github.com/pacifiic/augment-finetune-genomics.

## 1. Introduction

Deep learning models such as Basenji2 [1], Enformer [2], and Borzoi [3] have progressively improved in capturing long-range cis-regulatory interactions and predicting gene expression and chromatin accessibility. However, despite these advances, their predictive power remains limited when applied to personal genomes, where naturally occurring variants modulate expression in individual-specific ways. Recent studies [4–6] have shown that such sequence-based models exhibit limited predictive power across individuals, whereas PrediXcan [7], a statistical model trained on GTEx eQTL data, achieves substantially higher accuracy in predicting personal gene-expression levels. This discrepancy arises because current deep learning models are pre-trained exclusively on the reference genome, learning general regulatory syntax but lacking exposure to the population-level genetic diversity necessary for modeling personal transcriptomes.

Several studies have fine-tuned Enformer-based models on paired genome-transcriptome data to capture individual-specific variation, demonstrating that personal sequence diversity improves cross-individual prediction [8, 9]. To address the same challenge, Ramprasad et al. [10] proposed a convolutional model that integrates functional annotations, demonstrating the benefit of biological priors for improving generalization.

However, the limited availability of paired genome–transcriptome data remains a fundamental constraint. Publicly available datasets with both whole-genome sequencing (WGS) and RNA-seq measurements per individual are typically restricted to only a few hundred samples (e.g., GEUVADIS [11]), and even larger resources such as GTEx include fewer than one thousand individuals with matched WGS [12]. Moreover, access to high-quality paired datasets is often restricted due to privacy and data-sharing limitations. Given these constraints, pursuing data augmentation with synthetic genome–expression pairs represents a natural and necessary direction for improving cross-individual generalization.

To address this bottleneck, we propose a scalable fine-tuning framework that integrates simulated genomic diversity with GTEx-derived pseudo-expression labels. We first generate synthetic individual genomes using the `sim1000G` framework [13], which simulates realistic recombination and mutation events. Each simulated genome is assigned pseudo-expression values predicted by the PrediXcan model trained on GTEx eQTL data. [14, 15] Because the pseudo-labeled expressions are not on the same quantitative scale as true RNA-seq expression, we adopt a Bradley–Terry pairwise ranking objective [16] that learns relative expression rankings rather than absolute magnitudes. This preference-based approach allows for effective use of pseudo-labeled data while preserving the relative expression patterns observed in real data.

In our experiments, integrating both real and simulated data during fine-tuning yielded higher predictive accuracy than training on real data alone, highlighting the benefit of leveraging synthetic genetic diversity and pseudo-labeled expressions. Prior work evaluated deep learning models on the same variant-input, gene-level prediction task as Elastic Net [7], but reported only marginal gains—0.35% (from 0.283 to 0.284) in Rastogi et al. [8] and 0.37% in Ram-prasad et al. [10]. Ramprasad et al. further reported a 0.57% improvement from functional annotations; when combined multiplicatively, the two gains would amount to less than 1% overall improvement. Under an equivalent evaluation setting, our fine-tuned framework consistently outperformed PrediXcan, achieving a mean gene-level Pearson correlation improvement of 3.5% (from 0.311 to 0.322), showing a performance gain higher than those reported in previous studies.

**Fig. 1:**
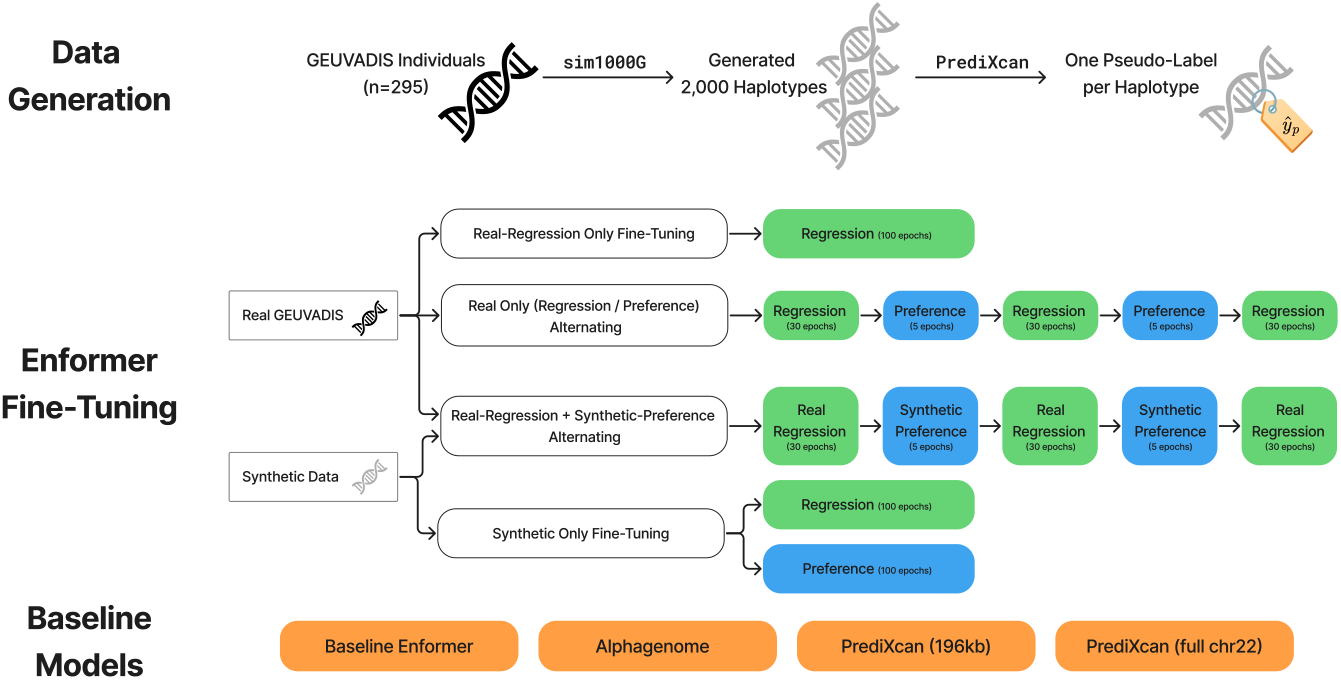
Overview of the data-generation and Enformer fine-tuning pipeline. `sim1000G` was used to generate synthetic haplotypes, which were pseudo-labeled by PrediXcan. The synthetic haplotypes were integrated with real GEUVADIS individuals to support a comprehensive set of fine-tuning configurations. Specifically, we evaluated: (i) real-only regression; (ii) real-only alternating regression–preference training; (iii) real–synthetic alternating schedules, which interleave real-data regression with synthetic-data preference optimization; and (iv) synthetic-only training, using either regression or preference-based objectives. Baseline comparisons include Enformer, AlphaGenome, and PrediXcan.

In summary, our study addresses key challenges in personalized transcriptome prediction by (1) synthesizing population-level genomic diversity and leveraging eQTL-based prediction models trained on large GTEx cohorts, and (2) developing an effective fine-tuning strategy that jointly integrates synthetic and real data.

## 2 Methods

### 2.1 Data Preprocessing

Genotype and RNA-seq expression data for 421 phased individuals were obtained from the GEUVADIS cohort [11], with genotypes originating from the 1000 Genomes Project [17]. The dataset was randomly split into 295, 42, and 84 individuals for training, validation, and testing, respectively. From the genes analyzed in Huang et al. [4], we selected those located on chromosome 22. For each gene, a 196,608 bp sequence centered on the transcription start site (TSS; ±98,304 bp) was extracted from the hg19 reference genome. After excluding genes overlapping ambiguous N regions in hg19, 42 genes were retained for all experiments.

To increase the genetic diversity of training samples, we simulated an additional 1,000 synthetic individuals using the `sim1000G` framework, which reproduces population-level recombination and linkage patterns. Predicted gene-expression levels for these simulated genomes were generated using PrediXcan’s eQTL models trained on GTEx v8. The resulting pseudo-labeled (sequence, expression) pairs were combined with real data for model fine-tuning. To prevent data leakage, synthetic genomes were simulated using only variants present in the training-set haplotypes, ensuring that no variants exclusive to the validation or test sets were included during simulation or training.

### 2.2 Model Architecture and Fine-Tuning

Our model builds upon the Enformer architecture, which predicts gene expression from long genomic sequences using convolutional and transformer layers. To adapt Enformer for individual-level prediction, we froze the convolutional feature extractor and applied parameter-efficient fine-tuning to the transformer component. Each input sequence was processed once through Enformer’s convolutional tower to obtain fixed-length feature vectors, which were then used as inputs during fine-tuning for computational efficiency.

This design was conceptually inspired by the parameter-efficient transfer learning framework proposed by Yuan et al. [18], which demonstrated the effectiveness of Low-Rank Adaptation (LoRA) [19] for fine-tuning large regulatory sequence models. Following this principle, we integrated LoRA modules into the self-attention mechanism of Enformer’s transformer layers, enabling targeted adaptation of attention projections without updating the full model.

To capture haplotype-specific regulatory effects, we adopted a dual-Enformer ensemble configuration that models maternal and paternal haplotypes independently. Two Enformer instances, denoted as Enformer_1_ and Enformer_2_, were jointly trained on paired haplotype sequences. To ensure symmetry and robustness, both input orderings—(Enformer_1_, Enformer_2_) = (maternal, paternal) and vice versa—were included during training. For synthetic sequences, where pseudo-expression labels were generated per haplotype rather than per individual, the same haplotype sequence was fed into both Enformer_1_ and Enformer_2_ to maintain architectural consistency. The outputs from the two Enformer instances were averaged to produce the final gene-expression estimate, which was optimized against either observed values or pairwise preferences between pseudo-labeled samples. During evaluation, the final prediction was computed as the mean of the two input-order configurations.

### 2.3 Training Objectives

We employed two fine-tuning objectives: SMAPE loss for regression-based training, as established in prior work [8], and a Bradley–Terry (BT) preference-based objective for training on mixed real and synthetic data. SMAPE minimizes the discrepancy between predicted and observed expression values for real WGS– RNA-seq pairs. In contrast, pseudo-expression values produced by PrediXcan are continuous outputs of a linear prediction model and therefore differ substantially in scale and distribution from true RNA-seq measurements. These scale mismatches make direct regression unreliable when real and synthetic samples are jointly used for supervision.

Rastogi et al. [8] introduced a ranking-based objective for fine-tuning gene-expression models, but their formulation relies on assumptions about the underlying count-generating process (e.g., Poisson/Skellum structure) that do not hold for PrediXcan-derived pseudo-labels. To provide a distribution-free and scale-invariant alternative, we adopt a Bradley–Terry preference model, which depends only on the relative ordering between samples and is therefore inherently robust to cross-domain scale differences.

Given a pair of samples (*A, B*), the model is trained to assign a higher predicted expression to the sample with the larger target value:

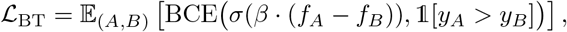

where *f*_*A*_ and *f*_*B*_ denote the predicted expression values, and *β* controls the sharpness of pairwise comparisons. The logistic sigmoid *σ*(·) maps the score difference to the probability that sample *A* should outrank sample *B*. Here, BCE denotes the binary cross-entropy loss. To account for gene-specific variability, *β* is scaled adaptively per gene based on the empirical distribution of expression differences.

### 2.4 Training Configuration

Model fine-tuning was performed for 100 epochs using four NVIDIA H100 GPUs under DeepSpeed ZeRO Stage 1 optimization [20]. A per-GPU batch size of 1 and a gradient accumulation step of 4 were used, resulting in an effective global batch size of 16. All transformer layers were fine-tuned using the AdamW optimizer (learning rate 1 × 10^−4^, weight decay 1 × 10^−3^), with a cosine learning rate scheduler including 1,000 warm-up steps, following prior work [8]. Mixed-precision training was conducted using the `bf16` format, and gradient clipping (maximum norm = 1.0) was applied to prevent gradient explosion. During preference-based training, pair sampling was controlled using an epoch- and rank-dependent seeding scheme to balance randomness and sample diversity across distributed workers. In the synthetic data integration experiments, the total number of training epochs and samples per epoch were kept identical to those in the real-data experiments. This design eliminated potential confounding effects from differing training steps, ensuring that performance improvements reflected the effect of data augmentation itself.

### 2.5 Evaluation Metrics

Model performance was evaluated following the same protocol as prior works [4, 8]. For each gene, we computed the Pearson and Spearman correlation coefficients between predicted and observed expression values across individuals in the same genes. Final performance metrics were obtained by averaging the correlations across all evaluated genes.

Model selection was based on validation performance: the checkpoint achieving the highest mean Pearson correlation across genes in the validation set was chosen and subsequently evaluated on the held-out test set. For the test set, both Pearson and Spearman correlations were computed in the same manner.

## 3 Experiments and Results

### 3.1 Experimental Overview

We conducted three groups of experiments to systematically evaluate the effectiveness of our proposed fine-tuning framework for personalized gene-expression prediction. First, we compared the model against three established baselines: the baseline Enformer model without fine-tuning, AlphaGenome [21], and PrediXcan. Second, we examined the predictive capacity of models trained solely on synthetic data, using either a regression-based objective or a preference-based objective. Finally, we investigated whether alternating real and synthetic fine-tuning phases could further enhance cross-individual generalization by interleaving regression and preference learning objectives across epochs.

### 3.2 Baseline Models

We first compared our fine-tuning framework with three baseline approaches for personalized gene-expression prediction: (i) the baseline Enformer model, (ii) AlphaGenome, and (iii) PrediXcan. All models were evaluated on the same set of 42 chromosome 22 genes across 84 GEUVADIS test individuals. Performance was measured using mean Pearson and Spearman correlation coefficients between predicted and observed expression levels across individuals.

As shown in Table 1, the baseline Enformer model achieved a mean Pearson correlation of 0.064 and a mean Spearman correlation of 0.047, confirming its limited generalization when applied to personal genomes. AlphaGenome, a follow-up study to Enformer that extends the input window to 1 Mb around each gene, improved performance substantially, with mean Pearson and Spearman correlations of 0.211 and 0.203 on the total RNA-seq track. Results on the polyA track were nearly identical (0.210 and 0.205).

**Table 1:** Baseline performance comparison on the same 42 genes, evaluated on 84 GEUVADIS test individuals. For each gene, Pearson and Spearman correlations were computed between predicted and observed expression values across individuals, and the reported metrics correspond to the mean correlations across the 42 genes. AlphaGenome results are provided for both polyA and total RNA-seq tracks.

| Model | Sequence Length | Mean Pearson | Mean Spearman |
| --- | --- | --- | --- |
| Enformer | 196 kb | 0.064 | 0.047 |
| AlphaGenome (polyA) | 1 Mb | 0.210 | 0.205 |
| AlphaGenome (total) | 1 Mb | 0.211 | 0.203 |
| PrediXcan | 196 kb | 0.299 | 0.289 |
| PrediXcan | full chr22 | 0.311 | 0.288 |

PrediXcan, an eQTL-based statistical model trained on GTEx v8 lymphoblas-toid cell line (LCL) data, achieved substantially higher performance than either deep learning baseline (mean Pearson 0.311, Spearman 0.288). This contrast highlights the gap between sequence-based models pretrained on a single reference genome and statistical predictors explicitly leveraging population-level eQTL information.

When restricted to a 196 kb genomic window centered on the transcription start site (TSS), the performance of PrediXcan decreased to Pearson 0.299 and Spearman 0.289. This reduction suggests that population-level SNP models also benefit from broader genomic context, implying the presence of regulatory eQTLs located outside Enformer’s 196 kb window.

### 3.3 Synthetic-Only Fine-Tuning

The purpose of this experiment is to evaluate the effectiveness of the augmented dataset itself. Enformer was fine-tuned solely on the pseudo-labeled synthetic data, using two independent models trained for 100 epochs each: one with the regression-based SMAPE loss and the other with the BT preference-based objective. Because pseudo-expression labels were generated at the haplotype level rather than the individual level, each synthetic training sample corresponded to a single haplotype sequence.

As shown in Table 2, the regression-based model trained on pseudo-labeled data exhibited limited predictive capacity (mean Pearson 0.171, Spearman 0.157), likely due to scale discrepancies between PrediXcan-derived expression values and real RNA-seq measurements. In contrast, the model optimized with the BT preference-based objective achieved substantially higher correlations (mean Pearson 0.307, Spearman 0.298), indicating that pairwise preference learning is more robust to absolute scale differences across pseudo-labeled samples. These findings suggest that although pseudo-labeled data may not reproduce absolute expression magnitudes, they still encode meaningful relative ordering information that can be effectively leveraged through ranking-based optimization.

**Table 2:** Performance of synthetic-only fine-tuning experiments on the same 42 genes, evaluated on 84 GEUVADIS test individuals. For each gene, Pearson and Spearman correlations were computed across individuals, and the reported values correspond to the mean correlations across the 42 genes. Results are shown for random seed 42.

| Objective | Mean Pearson | Mean Spearman |
| --- | --- | --- |
| Regression (SMAPE) | 0.171 | 0.157 |
| Preference-based (BT) | **0.307** | **0.298** |

### 3.4 Evaluating the Effect of Combining Real and Augmented Data on Fine-Tuning Performance

Having confirmed the standalone utility of the augmented (synthetic) dataset in the previous experiment, we next examined whether integrating real and synthetic fine-tuning phases could further improve predictive performance across individuals. To isolate the contribution of data augmentation from mere scheduling effects, we compare alternating real–synthetic schedules against a real-only regression baseline (100 epochs) and include a control schedule that uses *real* BT preference optimization instead of *synthetic* BT preference optimization.

In standard post-training pipelines for large language models (LLMs), supervised fine-tuning (SFT) and preference learning (e.g., DPO) are often performed sequentially, which is known to be suboptimal due to forgetting of supervised knowledge during later preference learning stages. Recent work has addressed this by introducing joint or alternating optimization; notably, *Fernando et al*. [22] showed that alternating SFT and DPO phases mitigates forgetting relative to purely sequential post-training. Motivated by this insight, we alternate between *real* regression and *synthetic* preference-based fine-tuning phases in the genomics setting.

#### Training schedules

We designed three training schedules under an equivalent computational budget, ensuring the same number of epochs and an equal amount of data processed per epoch: *(a)* real-only regression for 100 epochs; *(b)* alternating real regression and **synthetic** preference (30*R*_*r*_–5*S*_*p*_–30*R*_*r*_–5*S*_*p*_–30*R*_*r*_); and *(c)* alternating real regression and **real** preference (30*R*_*r*_–5*R*_*p*_–30*R*_*r*_–5*R*_*p*_– 30*R*_*r*_). All results below are reported as mean ± std over three fixed random seeds.

#### Results and interpretation

Table 3 shows that alternating regression and preference phases improves generalization over the real-only baseline. The real–synthetic alternating schedule (30*R*_*r*_–5*S*_*p*_–30*R*_*r*_–5*S*_*p*_–30*R*_*r*_) achieved superior performance compared to the other two fine-tuning strategies across both Pearson and Spearman correlation metrics. Because the 30*R*_*r*_–5*R*_*p*_ schedule maintains the same alternation pattern while using only real data, the additional gains observed in 30*R*_*r*_–5*S*_*p*_ are attributed not to scheduling but to the complementary signals from pseudo-labeled synthetic data, which enhance generalization by exposing the model to a broader distribution of individual-level genotypes. Compared with AlphaGenome and PrediXcan, the proposed real–synthetic alternating framework achieved higher performance under the same cross-individual evaluation setting. Specifically, it yielded mean Pearson and Spearman correlation improvements of 3.5% and 8.3%, respectively, over PrediXcan. While mean-based metrics summarize overall performance, they can obscure gene-specific variability. To determine whether the observed improvements were driven by a few genes or were consistent across genes, we conducted a per-gene correlation analysis (Figure 2). The per-gene distributions show that our model achieved positive *Δ* Pearson correlations for approximately 57% of genes in both PrediXcan baselines (full chromosome 22 and 196,608 bp window), indicating consistent improvements across the two comparisons.

**Table 3:** Performance comparison of baseline models and alternating fine-tuning schedules on the same 42 genes, evaluated on 84 GEUVADIS test individuals. For each gene, Pearson and Spearman correlations were computed across individuals, then averaged across the 42 genes. Reported values are mean ± std over three random seeds (42, 47, 52).

| Model / Training Schedule | Mean Pearson | Mean Spearman |
| --- | --- | --- |
| **Baselines** |  |  |
| AlphaGenome (1 Mb, total) | 0.211 | 0.203 |
| PrediXcan (196 kb) | 0.299 | 0.289 |
| PrediXcan (full chr22) | 0.311 | 0.288 |
| **Fine-tuning Schedules (ours)** |  |  |
| Real-only ($100R_r$) | $0.303 \pm 0.002$ | $0.295 \pm 0.003$ |
| $30R_r-5R_p-30R_r-5R_p-30R_r$ | $0.314 \pm 0.001$ | $0.306 \pm 0.002$ |
| $30R_r-5S_p-30R_r-5S_p-30R_r$ | **$0.322 \pm 0.004$** | **$0.312 \pm 0.005$** |

**Abbreviations:**
$R_r$ = real-data regression (real individuals, regression objective);
$R_p$ = real-data preference (real individuals, preference objective);
$S_p$ = synthetic-data preference (synthetic individuals, preference objective).
**Notation:** “ $30R_r-5S_p-\dots$ ” indicates 30 epochs of $R_r$ followed by 5 epochs of $S_p$ . Similarly, “ $100R_r$ ” denotes 100 epochs of regression-only fine-tuning on real individuals.

**Fig. 2:**
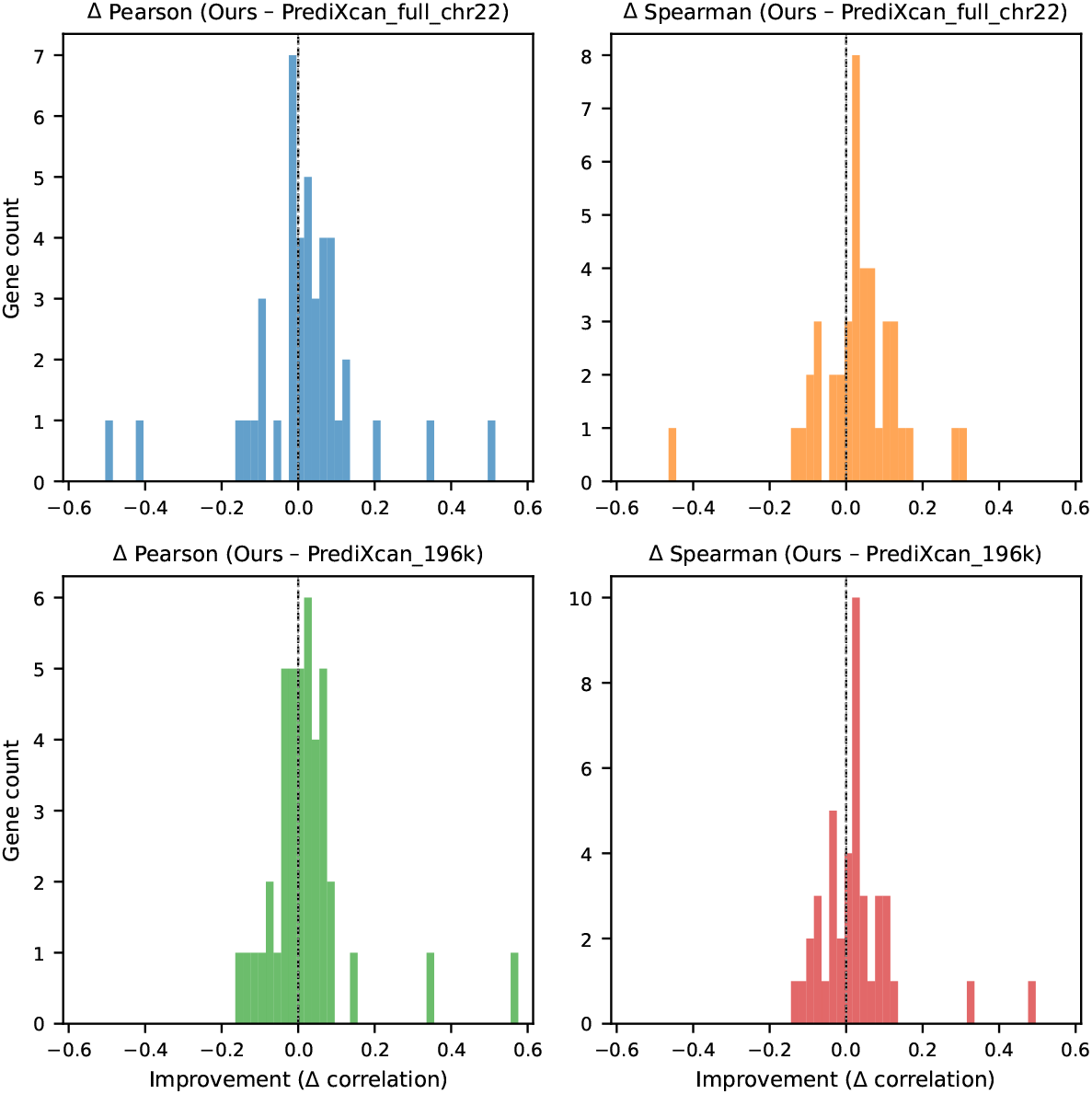
Per-gene improvement analysis comparing the proposed alternating fine-tuning schedule (**Ours**: 30*R*_*r*_–5*S*_*p*_–30*R*_*r*_–5*S*_*p*_–30*R*_*r*_) with two PrediXcan base-lines—one using the full chromosome 22 and another with 196,608 bp windows. Each bar indicates the correlation difference for an individual gene, where positive values denote higher Pearson or Spearman correlations achieved by our model. Results are from the run with random seed 42.

Notably, two specific genes, CRYBB2 and NDUFA6, showed markedly higher Pearson correlations under the full-length PrediXcan model, as observed in Figure 2 (comparing Ours with PrediXcan_full_chr22). This likely reflects the ability of broader genomic windows to capture distal regulatory signals beyond the 196 k bp receptive field of Enformer. For these genes, the differences in Pearson correlation (Ours - PrediXcan) were −0.497 and −0.422 under the full-chromosome comparison but became +0.069 and +0.151 in the 196 k bp window setting, suggesting that meaningful regulatory variants (eQTLs) likely reside beyond Enformer’s 196 k bp receptive field. PrediXcan, which leverages population-level eQTL weights without being constrained by fixed sequence windows, thus gains a clear advantage in capturing such distal regulatory effects.

## 4 Conclusions

We proposed a scalable fine-tuning framework that integrates simulated genomic diversity and eQTL-based pseudo-expression labels to improve personalized gene-expression prediction. By combining PrediXcan-derived pseudo labels with preference-based fine-tuning, the model effectively learns from relative expression patterns across simulated samples. First, even when trained solely on pseudo-labeled synthetic samples, the model achieved substantial predictive power, indicating that simulated genomes contain meaningful regulatory information that can be effectively exploited through ranking-based learning. Next, the alternating regression–ranking schedule surpassed the regression-only baseline, indicating that incorporating preference-based updates provides additional predictive benefit. Finally, the fine-tuning strategy that alternates between real and synthetic data consistently achieved the best performance across all settings.

Together, these results indicate that our pseudo-labeled synthetic genomes provide stable and meaningful data augmentation that enhances performance for personal gene-expression prediction.

